# Arteries are finely tuned thermosensors regulating myogenic tone and blood flow

**DOI:** 10.1101/2023.03.22.532099

**Authors:** Thieu X. Phan, Niaz Sahibzada, Gerard P. Ahern

## Abstract

In response to changing blood pressure, arteries adjust their caliber to control perfusion. This vital autoregulatory property, termed vascular myogenic tone, stabilizes downstream capillary pressure. We discovered that tissue temperature critically determines myogenic tone. Heating steeply activates tone in skeletal muscle, gut, brain and skin arteries with temperature coefficients (*Q*_10_) of ∼11-20. Further, arterial thermosensitivity is tuned to resting tissue temperatures, making myogenic tone sensitive to small thermal fluctuations. Interestingly, temperature and intraluminal pressure are sensed largely independently and integrated to trigger myogenic tone. We show that TRPV1 and TRPM4 mediate heat-induced tone in skeletal muscle arteries. Variations in tissue temperature are known to alter vascular conductance; remarkably, thermosensitive tone counterbalances this effect, thus protecting capillary integrity and fluid balance. In conclusion, thermosensitive myogenic tone is a fundamental homeostatic mechanism regulating tissue perfusion.

**One-Sentence Summary:** Arterial blood pressure and temperature are integrated via thermosensitve ion channels to produce myogenic tone.

## Introduction

The cardiovascular system has critical roles both in transporting oxygen/nutrients and in thermoregulation. To achieve these goals the cardiac output must be appropriately distributed among tissues. Tissue blood flow is regulated by the autonomic nervous system, humoral transmitters, shear stress and metabolic factors (*1, 2*). In addition, small arteries and arterioles possess the intrinsic capacity to constrict or dilate in response to changes in intravascular pressure. This autoregulatory mechanism is known as the vascular myogenic response or myogenic tone (*3, 4*). Vascular myogenic tone (*1, 2*), enables a highly localized, fine-tuned control of blood perfusion and protects against fluctuations in capillary pressure that would negatively affect fluid transport or compromise vessel integrity (*3*). Indeed, a diminished myogenic tone accompanies diabetes, obesity (*4*), coronary artery disease (*5*), hypertrophic cardiomyopathy (*6*) and ageing (*7, 8*); all diseases or states with impaired hemodynamic regulation.

For over 120 years, blood pressure has been considered the primary determinant of myogenic tone (*1, 9*). Specifically, blood pressure exerts a circumferential stretch of the surrounding vascular smooth muscle cells (VSMC) (*9*) leading to membrane depolarization, the activation of L-type Ca^2+^ channels and a subsequent increase in myoplasmic [Ca^2+^] necessary to trigger contraction (*10-13*). Recent studies have identified G-protein coupled receptors (GPCR) as key VSMC mechanosensors (*9, 14*). In turn, these GPCRs activate transduction channels, which mediate membrane depolarization. Putative transduction channels include members of the transient receptor potential (TRP) family (*16-20*) and ANO1 (*21*).

Temperature prominently affects blood flow both indirectly (via the autonomic/sensory nervous systems) and directly. Raised local temperature directly increases blood flow and this effect has been demonstrated in diverse tissues including skeletal muscle, bone, gut, and lung (*15-21*). Heat-induced hyperemia can be explained by changes in blood rheology (such as increased blood fluidity, RBC deformability and dispersion) (*22, 23*) and/or vasodilation (*19*). These direct effects of temperature on blood flow represent an often, overlooked consideration in the circulatory system, which carries blood through tissues of varying temperatures. Indeed, the skin and skeletal muscle in the body’s thermal shell experience large temperature changes that at distal sites may exceed 10-15°C (*15, 24, 25*). Further, core body temperatures display daily variations that are amplified by exercise, thermal stress and metabolism, including significant deviations during torpor (*26*). If unchecked, these fluctuating tissue temperatures would lead to large changes in tissue perfusion.

In this study, we show that arteries are intrinsically, highly thermosensitive and generate myogenic tone by integrating both pressure and temperature stimuli. In this manner, the myogenic response optimally regulates blood flow under diverse physiological conditions.

## Results

### Myogenic tone is steeply heat-dependent and tuned to tissue temperature

Recently, our lab uncovered a role for TRPV1 in the myogenic tone of skeletal muscle. TRPV1 is a multimodal channel which responds to extracellular protons, phospholipase C (PLC) signaling, and heat (*27*). Intriguingly, several other candidate transduction channels for myogenic tone are thermosensitive, including TRPM4 (*28*) and ANO1 (*29*). We therefore hypothesized that myogenic tone is dynamically regulated by temperature. We first explored whether temperature specifically regulates tone in radial muscle branch arteries (**Fig. S1A**). In isolated, pressurized (80 mmHg) arteries, myogenic tone was mostly absent below ∼28°C, but developed steeply to a maximum of 45% as temperature was incrementally raised to 41°C (**Figs. 1A-D**). The mean temperature for half-maximal activation (T_1/2_), calculated by fitting data from individual arteries, was 33.4 ± 1.8 °C (n=11). Analysis of the temperature dependence using an Arrhenius plot, revealed a steep component with a threshold of ∼27-28°C and a mean temperature coefficient (*Q*_10_) of 12.6 ± 1.8 (n=6, **Fig. 1D and Fig. S2**). Further, rapid cooling from 37°C to 32°C abruptly dilated arteries (**Fig. 1C**), supporting a highly temperature-sensitive process.

**Figure 1.**
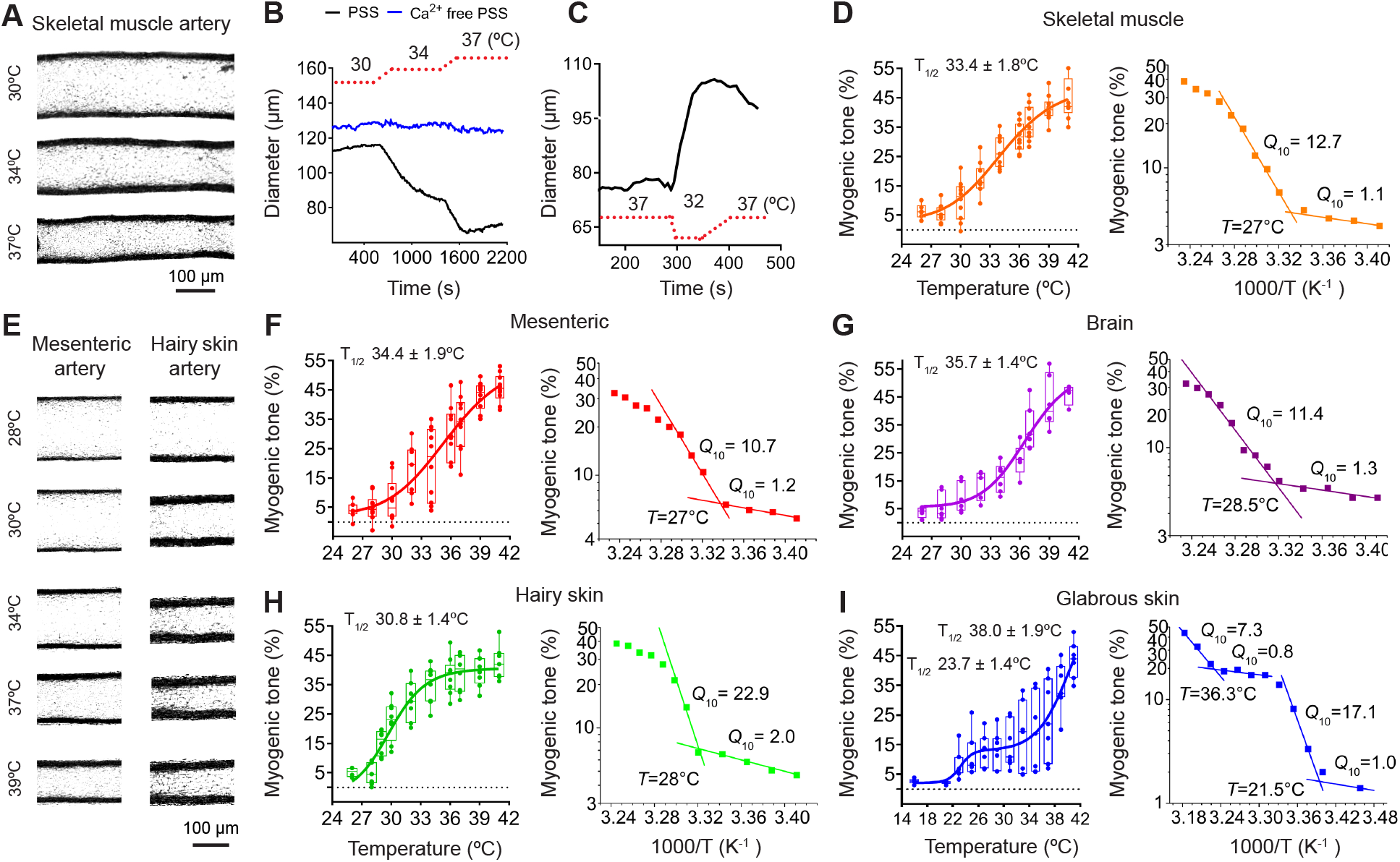
Myogenic tone is highly thermosensitive. **(A&E)** Representative photographs of pressurized (80 mmHg) skeletal muscle, mesenteric and hairy skin arterioles at different temperatures. **(B&C)** Changes in the diameter of pressurized (80 mmHg) skeletal muscle arteries in response to an increase or decrease in bath temperature (indicated by red dotted line). PSS: physiological salt solution **(D-I)** Mean myogenic tone (%) versus temperature and representative Arrhenius plots yielding indicated *Q*_10_ values and threshold temperatures (T), for skeletal muscle (n=11), mesenteric (n=11), pial (n=7), hairy skin (n=10) and glabrous skin (n=7) arteries. The mean half maximal temperature (T_1/2_) for tone was derived from fits to modified Hill equations of individual arteries.

Next, to test whether the dynamic temperature sensitivity of myogenic tone is a general property of resistance vessels, we studied pressurized arteries (80-150μM diameter) isolated from the gut (mesenteric), brain (pial) and skin (hairy and glabrous) (**Figs. S1B-E**). Remarkably, we found that myogenic tone in all these arteries is temperature-dependent over a narrow range; albeit the T_1/2_ was tissue-specific (**Figs. 1E-I**). In mesenteric arteries, heating activated a maximal tone of ∼46% with a T_1/2_ of 34.4 ± 1.9°C (n=11) and a *Q*_10_ of 11.7 ± 1.1 (n=5) **(Figs. 1E&F and Fig. S2)**. In pial arteries, heating activated 49% tone with a T_1/2_ of 35.7 ± 1.4°C (n=6) and a *Q*_10_ of 11.3 ± 1.3 (n=4) **(Fig. 1G and Fig. S2)**. In “hairy” skin arteries, heating activated 44% tone with a T_1/2_ of 30.8 ± 1.4°C (n=10) and a *Q*_10_ of 20.0 ± 2.4 (n=4) **(Figs. 1E&H and Fig. S2)**. In glabrous skin (digital) arteries, heating activated a bi-phasic response; an initial component of ∼13% tone with a T_1/2_ of 23.7 ± 1.4°C (n=7) and an additional component of 40% with a T_1/2_ of 38.0 ± 1.9°C **(Fig. 1I)**. The respective *Q*_10_ values were 17.3 ± 3.6 (20-26°C, n=5), and 11.9 ± 5.0 (>35°C, n=5) **(Fig. 1I and Fig. S2)**. Thus, a steep temperature dependence for myogenic tone appears to be a common property of all arteries. Further, these data explain the earlier observation that physiological temperatures are required to measure myogenic tone (*30*).

All the arteries we examined reside in tissues with distinct temperatures, reflecting their location in the body’s thermal core or shell (**Fig. 2A**). The temperature of the blood in small arteries (50-150μM) approximates the tissue temperature due to heat loss and counter-current heat exchange with veins (*31, 32*). Thus, VSMCs of small peripheral arteries sense the tissue temperature. Notably, the T_1/2_ for myogenic tone from individual arteries showed a high degree of correlation (r^2^=0.93, P<0.0001) with published tissue temperatures (*33-38*) (**Fig. 2B**). Thus, the thermosensitivity of myogenic tone (hereafter abbreviated as thermo-tone) is tuned to tissue temperature.

**Figure 2.**
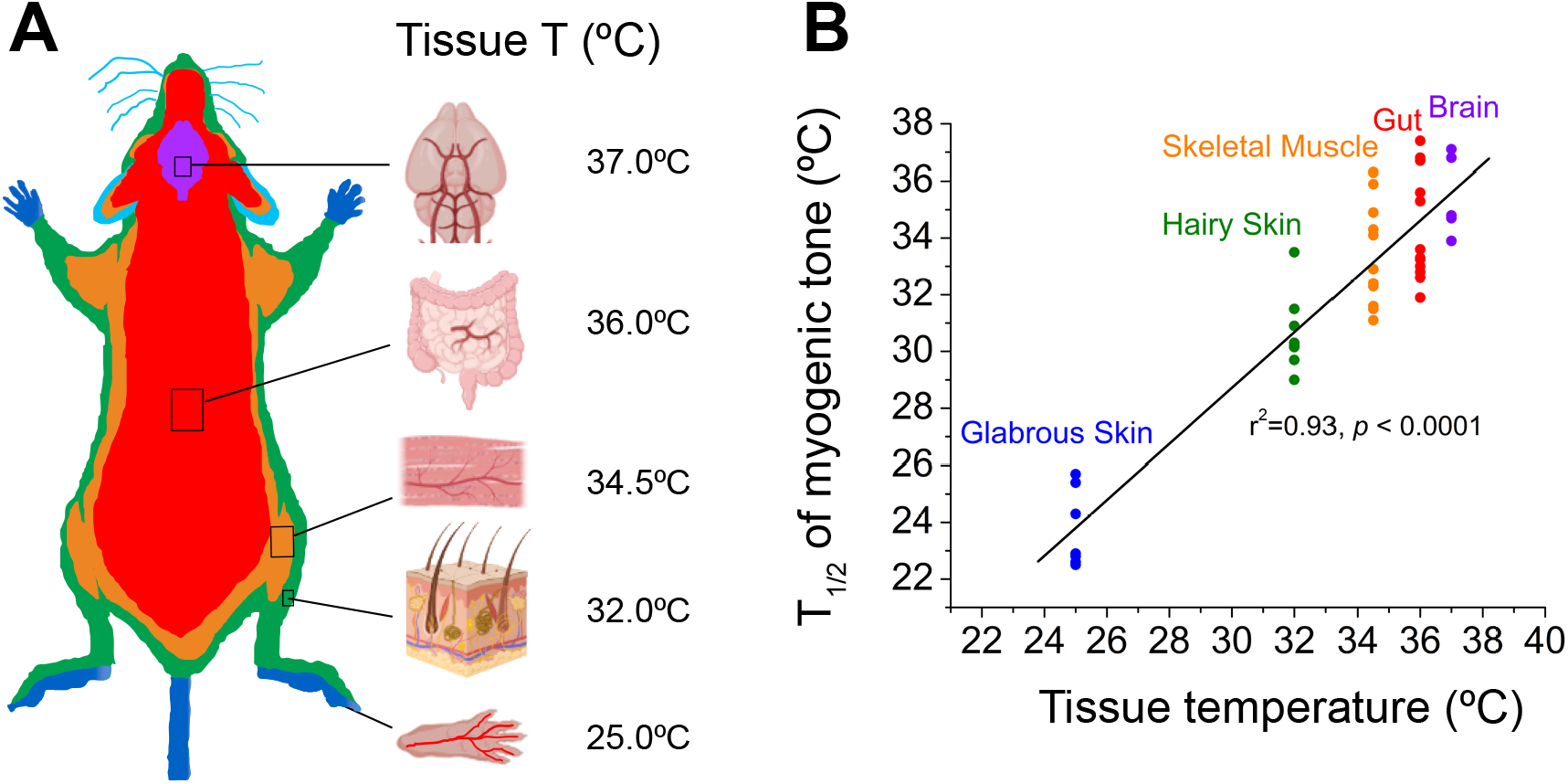
The thermosensitivity of myogenic tone correlates with tissue temperature. **(A)** Illustration of thermal tissue variations in mice. Approximate resting tissue temperatures were obtained from published values for skeletal muscle, brain, gut, hairy skin and glabrous skin (*33-38*) at an ambient T of 21-22°C. **(B)** Plot of myogenic tone T_1/2_ for individual arteries from skeletal muscle (n=11), mesenteric (n=11), pial (n=6), hairy skin (n=10) and glabrous skin (n=7) versus the tissue temperature.

### Pressure and temperature are independent sensors for myogenic tone

Next, we explored the underlying basis for thermo-tone. Temperature affects all proteins and enzymatic reactions with a *Q*_10_ of ∼2-3. In contrast, the *Q*_10_ for myogenic tone of ∼11-20 suggests a highly thermosensitive mechanism, consistent with a role for specialized heat-sensitive proteins.

To identify the locus for temperature sensitivity, we first tested the interrelationship between intraluminal pressure and temperature in skeletal muscle arteries. **Figures 3A&B** show the effect of varying the pressure (0-120 mmHg) at fixed temperatures. At 32°C, raising the intraluminal pressure evoked a maximal tone of 15% and the half-maximal response (P_1/2_) was obtained at 68 ± 14 mmHg. Increasing the temperature to 35°C and then 39°C significantly augmented the maximal tone to 25% and 40%, respectively, with only a modest reduction in the P_1/2_ to 58 ± 6 and 47 ± 12 mmHg respectively (**Fig. 3B**). Similarly, we measured the temperature sensitivity at different fixed pressures (40, 80 and 120 mmHg). **Figures 3C&D** show that raising the pressure increased the maximal tone without significantly altering the T_1/2_ of ∼34°C. Thus, both pressure and temperature are critical stimuli that are sensed largely independently yet integrated to generate myogenic tone. Further, these results indicate that mechanosensors in VSMCs do not significantly contribute to thermosensing.

**Figure 3.**
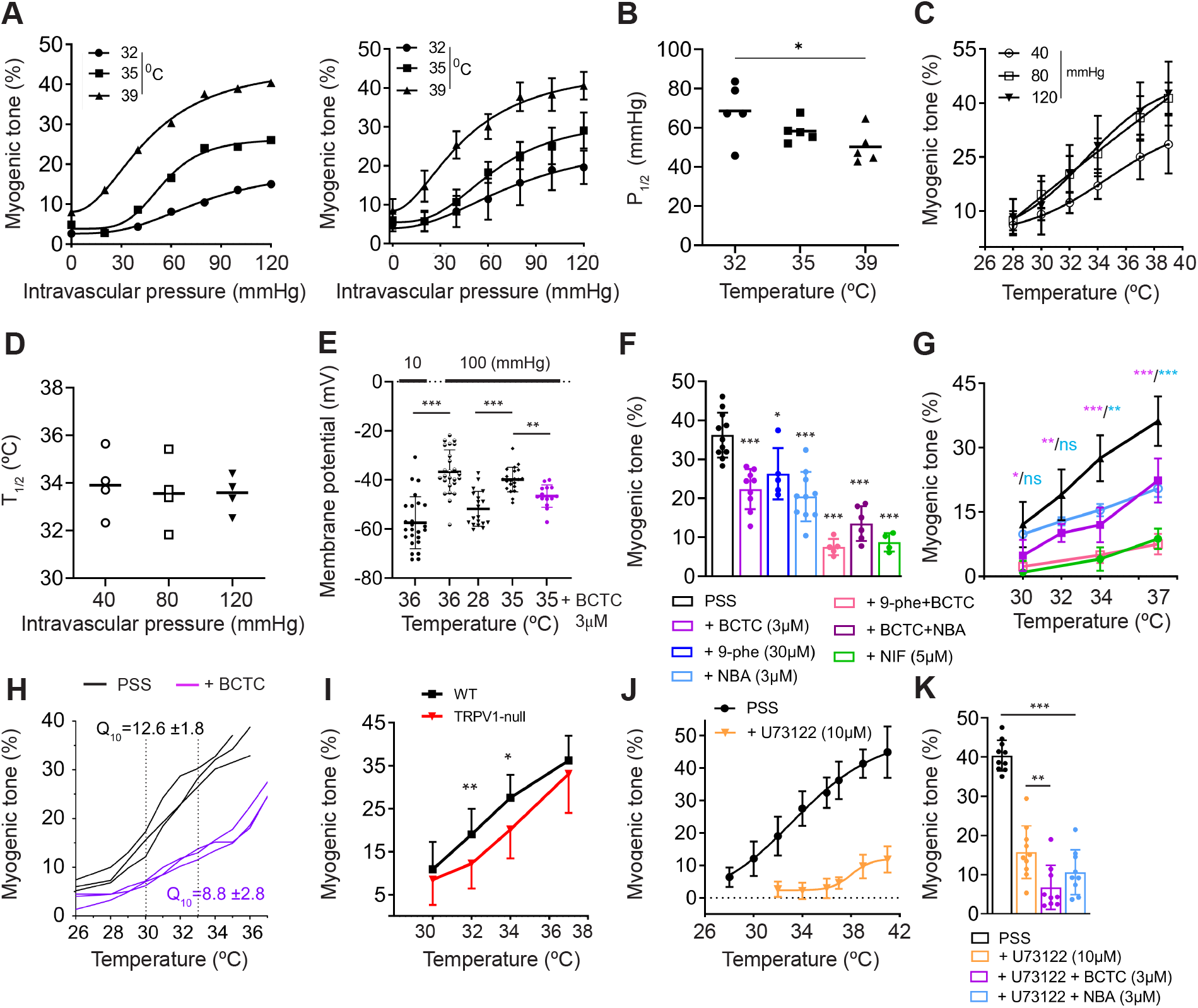
Distinct pressure and temperature sensors mediate myogenic tone in skeletal muscle. **(A)** Myogenic tone versus pressure in a single skeletal muscle arteriole at 32°C, 35°C and 39°C (*left*) and the mean responses of arteries from six animals (*right*). **(B)** Plot of half maximal pressure (P_1/2_) versus temperature (n=5, one-way ANOVA, *P<0.05). P_1/2_ values were obtained from the data in *a*. **(C&D)** Myogenic tone versus temperature, measured at intraluminal pressures of 40, 80 and 120 mmHg and plot of T_1/2_ versus pressure (n=4). **(E-K) TRPV1 and TRPM4 mediate thermo-tone**. (E) Membrane potential in smooth muscle cells of skeletal muscle arteries pressurized to 10 mmHg (n=25) or 100 mmHg (36°C, n=25; 28°C, n=19; 35°C, n=21; 35°C plus BCTC, n=14; one-way ANOVA, **P<0.01, ***P<0.001). (F) Summary of myogenic tone at 37°C and 80 mmHg under control conditions (PSS, n=9) or during pharmacological inhibition with the TRPV1 inhibitor, BCTC (3μM, n=4), TRPM4 inhibitors, 9-phenanthrol (9-phe, 30μM, n=5) and NBA (3μM, n=6), 9-phe+BCTC (n=5), NBA+BCTC (n=6) and L-type Ca^2+^ channel inhibitor nifedipine (NIF, 5 μM, n=4) (one-way ANOVA, *P<0.05, ***P<0.001). (G) Pharmacologic inhibition of thermo-tone at different temperatures (n=4, one-way ANOVA, *P<0.05, **P<0.01, ***P<0.001). (H) Plots of myogenic tone versus temperature for individual arteries under control conditions (PSS) or in the presence of BCTC. The mean *Q*_10_ derived from Arrhenius plots between 30-33°C are shown. (I) Temperature-dependent myogenic tone (80 mmHg) in WT (n=12) versus TRPV1-null (n=3) arteries (unpaired *t*-test, *P<0.05, **P<0.01). (J) Temperature-dependent myogenic tone (80 mmHg) in control arteries (PSS) (data re-plotted from Fig.1D, T_1/2_ =33.4°C), and arteries treated with the PLC inhibitor, U73122 (10μM, n=10). The T_1/2_ =39.1°C. (K) The maximal myogenic tone at 41°C for control (PSS, n=10), U73122 (n=10), U73122 + BCTC (n=9) and U73122 +NBA (n=9) (one-way ANOVA, ** P<0.01, ***P<0.001).

To understand how temperature and pressure are integrated, we tested for thermosensitivity in the signaling pathways downstream of pressure sensing. Previous studies have shown that stretch-mediated depolarization of VSMCs, followed by activation of L-type Ca^2+^ channels (Ca_v_1.2), accounts for most tone (*10-13*). Accordingly, we found that the Ca^2+^ channel blocker, nifedipine inhibited ∼80% of tone in skeletal muscle arteries (**Figs. 3F&G**). Heating had only a modest effect on the residual nifedipine-insensitive tone (**Fig. 3G**). This is consistent with the minimal temperature sensitivity of VSM contraction (*Q*_10_ of 1.8) (*39*). Further, the gating of L-type Ca^2+^ channels is only weakly temperature-dependent (*40*). Together, these data suggest that temperature sensing resides upstream of Ca^2+^ entry. Indeed, we found that heating regulated the stretch-induced membrane depolarization of VSMCs; at 28°C, pressurized cells remained hyperpolarized, but increasing the temperature to 36°C depolarized the membrane potential by 20 mV (**Fig. 3E**). Thus, these data suggest that the transduction channels that mediate membrane depolarization are the locus for thermo-tone.

### TRPV1 and TRPM4 mediate thermo-tone in skeletal muscle arteries

TRPV1 and TRPM4 are putative transduction channels for myogenic tone in skeletal muscle arteries, activated via a PLC-dependent pathway (*41*). To test a role for TRPV1 in thermo-tone, we used BCTC, an antagonist with demonstrated on-target selectivity for TRPV1 in VSMCs (*41*). BCTC inhibited VSMC membrane depolarization in pressurized arteries at 35°C (**Fig. 3E)**. Further, BCTC inhibited myogenic tone (80 mmHg) in a temperature dependent manner (**Figs. 3F-H**). BCTC had a greater inhibitory effect between 30°C to 34°C (∼60% block) compared with 37°C (∼40% block), supporting a larger relative contribution of TRPV1 to tone at these lower temperatures (**Fig. 3G)**. Accordingly, BCTC reduced the *Q*_10_ over the 30-33°C temperature range (**Fig. 3H)** and produced a right shift in the temperature-myogenic tone relationship (**Fig. S3)**; the T_1/2_ increased from 33.4 ± 1.8 °C (n=11) to 35.9 ± 0.6 (n=5, P=0.01). Thus, pharmacological inhibition of TRPV1 reduced the thermosensitivity of myogenic tone. Likewise, myogenic tone (80 mmHg) in TRPV1-deficient arteries was significantly depressed at 32°C and 34°C (**Fig. 3I**). However, tone at 37°C was normal in TRPV1-null arteries, suggesting compensation for the loss of TRPV1 at higher temperatures. Indeed, disruption of the TRPV1 gene leads to increased expression of TRPM4 (*41*). To test a role for TRPM4 in thermo-tone, we used the antagonist, 9-phenantrol (a commonly employed, yet non-selective drug), and a novel highly selective inhibitor, 4-Chloro-2-(1-naphthyloxyacetamido) benzoic acid (NBA). Both compounds significantly blocked myogenic tone by ∼40% at 37°C (**Fig. 3F)**. The inhibitory effect of NBA was reduced at lower temperatures (**Fig. 3G)**; unlike BCTC, NBA did not significantly inhibit tone at 30°C and 32°C, suggesting that TRPM4 is activated at a slightly higher temperature range. Further, combining NBA with BCTC suppressed tone to the same extent as blocking L-type Ca^2+^ channels, suggesting that TRPV1 and TRPM4 are the major contributors to thermo-tone in the 30-37°C temperature range (**Fig. 3F**). Thus, in skeletal muscle arteries, TRPV1 and TRPM4 appear to confer a largely overlapping thermal sensitivity of myogenic tone, with TRPV1 making a greater contribution to thermosensing at 30-34°C.

The activation of TRPV1 at 30°C in VSMCs may seem surprising, since thermal activation of TRPV1 typically occurs at greater than 40°C (*27*). However, allosteric modulators are known to substantially lower the heat threshold. Indeed, PLC-mediated signaling, leading to phosphoinositide depletion, generation of arachidonic acid metabolites or PKC activation, triggers activation of TRPV1 at <32°C (*42*) and this may be accompanied by a substantial reduction in the *Q*_10_ (*43*). Accordingly, we tested whether PLC regulates thermo-tone. **Figures 3J&K** show that the PLC inhibitor, U73122, reduced the maximal tone in pressurized arteries from ∼45% to 12% and shifted the temperature threshold from 28°C to 38°C and the T_1/2_ from 33.4°C to 39.1°C. Furthermore, the TRPV1 inhibitor, BCTC significantly inhibited the residual tone in U73122-treated arteries (P<0.001) (**Fig. 3K**). Thus, PLC signaling (with a major contribution of TRPV1) tunes skeletal muscle thermo-tone close to the resting temperature of skeletal muscle.

Next, to assess the role of thermo-tone *in vivo*, we exploited the well-described pharmacologic effects of TRPV1 antagonists on systemic blood pressure. When administered acutely to mice, TRPV1 inhibitors reduce the systemic blood pressure (*41*). This response is attributed to blockade of TRPV1-dependent myogenic tone; the effect persists after the deletion of TRPV1-positive sensory neurons and is therefore selectively mediated by vascular TRPV1 (*41*). We therefore measured the systemic blood pressure response to the TRPV1 antagonist, BCTC, at normothermic and hypothermic conditions (**Fig. 4A**), verified by measuring both the core and skeletal muscle temperatures (T_Core_/_Sk.M_). As previously reported, BCTC (3 mg/kg) administered to mice at a near normal T_Core_/_Sk.M_ of 36.4/33.0°C produced an immediate decrease in blood pressure of ∼10 mmHg (**Figs. 4B-E**). In contrast, when the T_Core_/_Sk.M_ was lowered to 32/29.0°C, BCTC failed to change the blood pressure (**Figs. 4B-E**). Hypothermia alone did not significantly affect the resting blood pressure (**Figs. 4B&D**) and therefore, the results are consistent with a loss of TRPV1-dependent thermo-tone. Further, intravital imaging showed that raising the local temperature surrounding the radial muscle branch arteries from 30°C to 37°C significantly decreased the vessel diameter (**Figs. 4F&G**). Thus, these data show that physiologic changes in temperature regulate myogenic tone *in vivo*.

**Figure 4.**
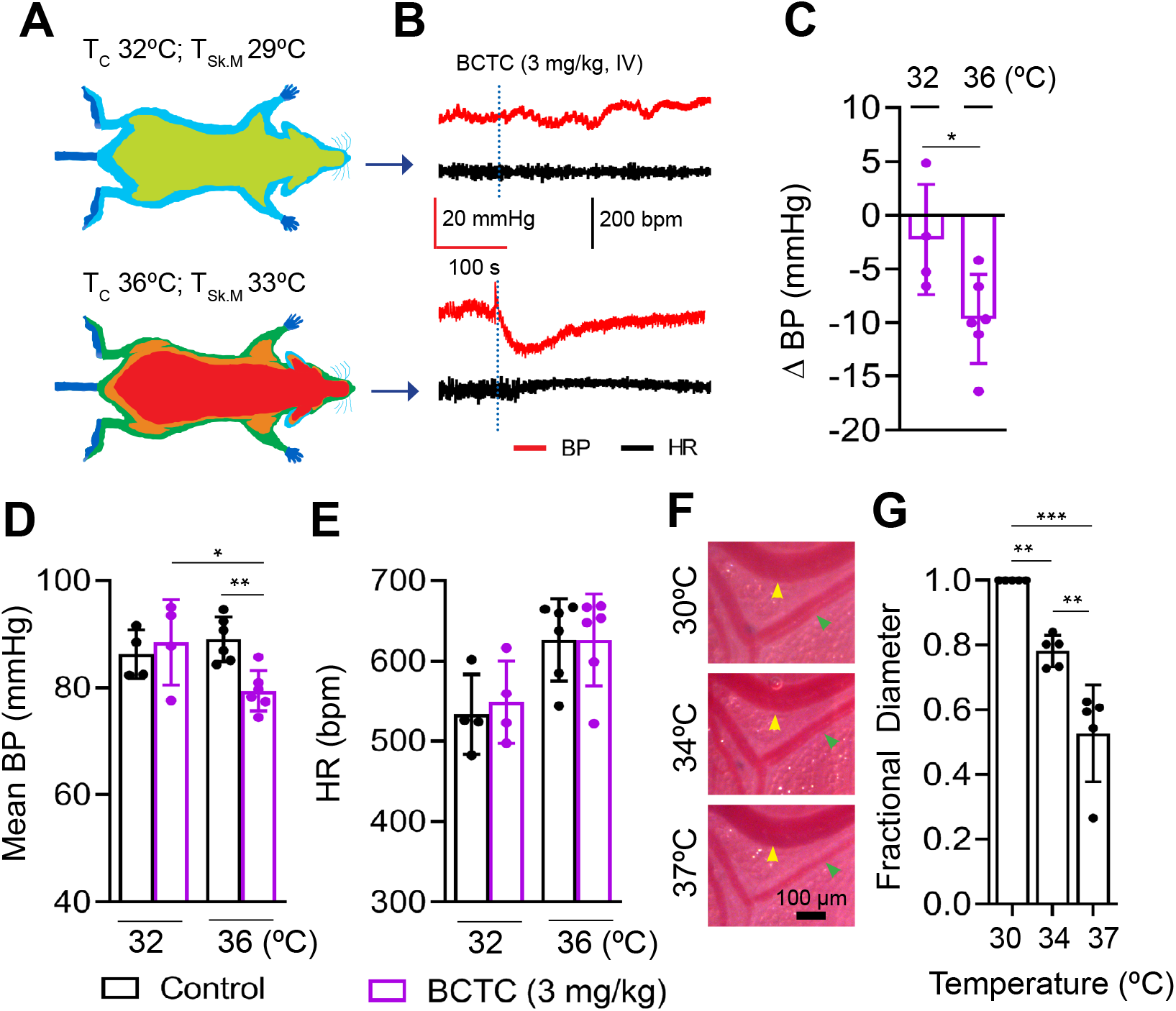
Temperature regulates myogenic tone *in vivo*. **(A)** Anesthetized mice were maintained at a core/skeletal muscle temperature (T_C_/T_Sk.M_) of 32°C/29°C or 36°C/33°C. **(B)** Representative systemic blood pressure (BP) and heart rate (HR) recording during i.v. administration of the TRPV1 inhibitor, BCTC (3mg/kg, dotted line) **(C)** The change in the mean BP evoked by BCTC at different T_C_ (32°C, n= 4; 36°C, n=6, unpaired *t*-test, *P<0.05). **(D&E)** Summary of mean arterial pressure and heart rate (32°C n= 4; 36°C, n=6; one-way ANOVA, *P<0.05, **P<0.01). **(F&G)** Intravital imaging of mouse radial muscle feed arteriole (green arrow) and veins (yellow arrow) shows arteriole constriction in response to local perfusion of heated PSS (30, 34 and 37°C, one-way ANOVA, n = 5, \*\**P* < 0.01, \*\*\**P* < 0.001).

### Thermo-tone counteracts heat-evoked changes in blood flow

Our results indicate that myogenic tone increases with temperature. This may seem counterintuitive since earlier studies have demonstrated that local tissue heating increases blood flow (*17-19*). Therefore, to understand how thermo-tone affects blood perfusion, we performed laser speckle imaging of the skeletal muscle radial branch arteries (**Figs. 5A&B**). In these experiments we varied the local tissue temperature while maintaining a stable core temperature of 37°C. Also, to prevent any confounding effects of changes in sympathetic tone, we included the α1-adrenergic antagonist, prazosin, in the super-perfusate. **Figures 5B-D** show that step increases in the local temperature from 28°C to 34°C and 39°C modestly increased perfusion (by 31% and 80%, respectively), in line with previous studies (*17, 18*). To test whether changes in myogenic tone regulate this heat-induced response, we blocked tone using either a Ca^2+^-free medium (supplemented with sodium nitroprusside) or nifedipine. After blocking myogenic tone, heating markedly increased perfusion (by 221% and 247%, respectively, **Figs. 5B-D)**. Remarkably, this represents a ∼25% increase in perfusion for each degree (over the 28-34°C range), Thus, inhibiting myogenic tone unmasks the full effects of temperature on blood flow. Similarly, treatment with BCTC plus NBA increased perfusion at 34°C and 39°C but not 28°C, consistent with the relaxation of TRPV1/TRPM4-dependent thermo-tone (**Figs. 5B-D)**. Taken together, these data show that thermo-tone counteracts the temperature-induced facilitation of blood perfusion.

**Figure 5.**
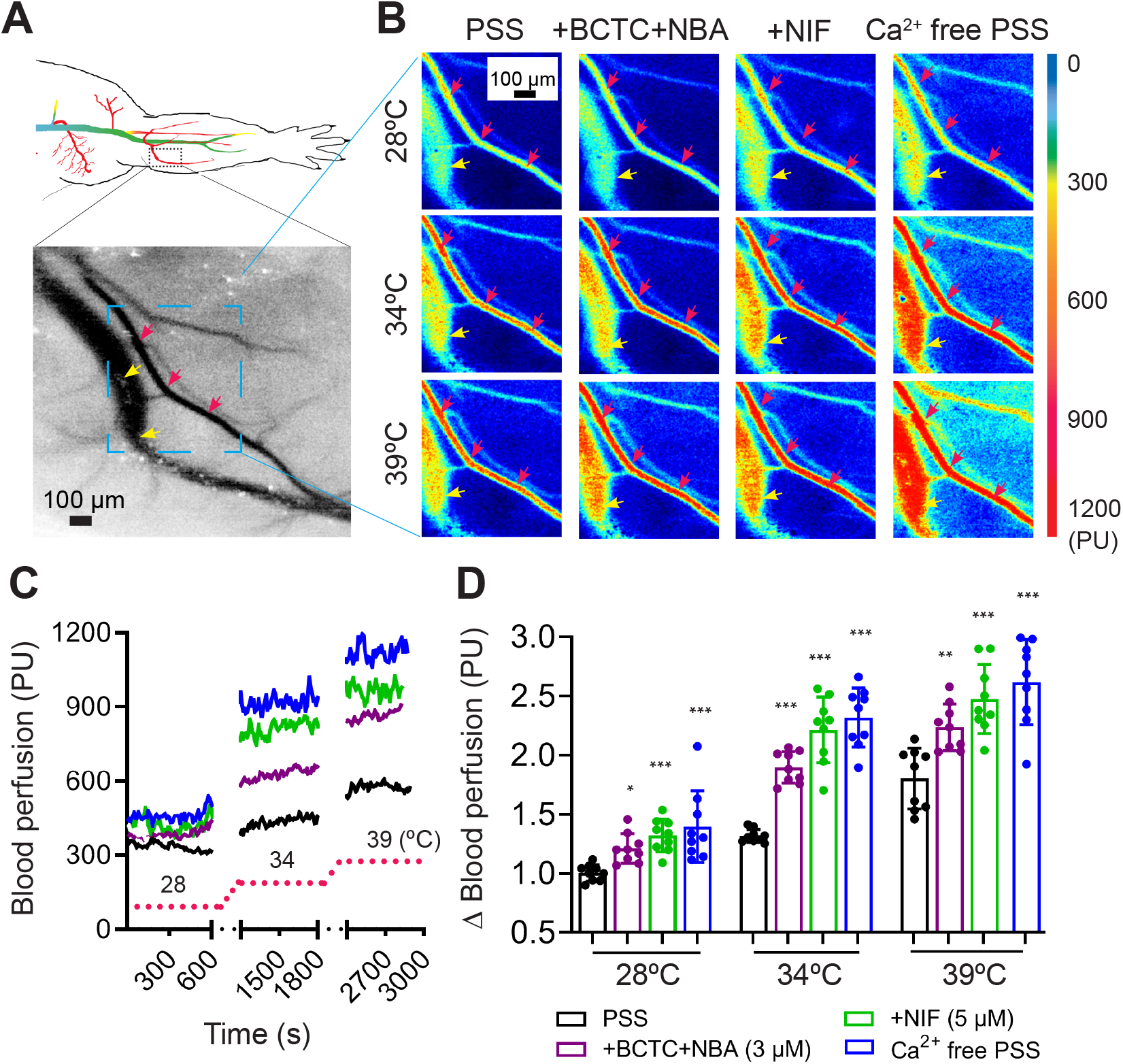
Thermo-tone opposes heat-evoked increases in blood flow. **(A)** *Top*, illustration of the intravital imaging field (dotted box) in the mouse forearm. *Bottom*, expanded view showing the radial muscle branch artery (red arrows) and a vein (yellow arrows). **(B)** Representative laser speckle contract images and **(C)** Relative blood perfusion during limb heating from 28 to 39°C with physiological salt solution (PSS, black), BCTC+NBA (purple), nifedipine (NIF, green) and Ca^2+^ free/ sodium nitroprusside PSS (blue). **(D)** Mean changes in blood perfusion (20°C, n=5; 28-39°C, n=9, one-way ANOVA, *P<0.05, **P<0.01, ***P<0.001). Note: all media contained the α1 adrenergic inhibitor, prazosin (10 μM).

## Discussion

Our results show that myogenic tone is steeply temperature dependent and tuned to tissue temperatures. Thus, arterial caliber and blood flow are maximally sensitive to small variations from the tissue’s resting temperature. Blood flow depends on the 4^th^ power of the vessel radius, therefore in principle, a one-degree °C change in temperature would generate up to a 40% change in blood flow in the most temperature sensitive arteries. While core body temperature is generally maintained as the ambient temperature changes, the skin experiences constant temperature fluctuations. Fittingly, we demonstrate that myogenic tone in skin arteries exhibits the greatest temperature sensitivity, with a *Q*_10_ of ∼20. The glabrous skin covers heat-exchange organs (*e*.*g*., the human hand or mouse paw) and thus has two distinct resting temperatures based on the level of sympathetic tone (*25*). The first state, which occurs at ambient temperature, operates under high resting sympathetic tone with minimal blood flow and a skin temperature just above the ambient level. Second, when there is little to no sympathetic tone (when overheated), glabrous skin temperature more closely matches core temperature as heated blood from the core flows to the organ (increase flow of >500%) to dump heat into the environment. The biphasic response to temperature in glabrous skin (**Fig. 1I)** matches these two distinct states.

Our analysis shows that the average maximum myogenic tone in skeletal muscle, gut, brain, and skin at 39-41°C is between 44% and 53%. In contrast, previous studies using fixed temperatures report varying levels (from ∼15-45%) of myogenic tone in defined mesenteric and cerebral arteries (*44-50*). One obvious explanation for the reported variability is that the temperature conditions employed in some studies were insufficient to fully activate tone. This suggests that the maximum myogenic tone does not vary drastically between tissues, rather that the thermosensitivity of myogenic tone is tissue dependent.

At first glance, thermo-tone may appear to contradict key features of thermal physiology. However, myogenic tone is just one component of hemodynamic regulation, acting in combination with metabolic, humoral, and neurogenic factors. Thus, whole body/skin heating triggers a sympathetic nerve-mediated increase in cutaneous blood flow (*17, 51*) and this effect can more than compensate for the effects of thermo-tone. Similarly, during exercise, the temperature of skeletal muscle can increase significantly and the resultant increase in thermo-tone would appear to imperil the required increase in blood supply. However, metabolic factors produced by contracting muscle are well known to inhibit myogenic tone, such that there is an ascending relaxation of the supply vessels (*52*). Indeed, in this context, increased thermo-tone may be beneficial by re-directing blood flow to actively contracting muscle units.

Our findings suggest that VSMCs integrate the output of distinct pressure and temperature sensors to generate myogenic tone (**Figs. 3A-D)**. Although the precise molecular mechanisms underlying the myogenic response are still unresolved, there is strong evidence supporting a role for GPCRs as mechanosensors. Indeed, the Angiotensin II Receptor AT1 plays a key role in the myogenic tone of mesenteric, brain, and skeletal muscle arteries (*9, 47-49, 53*). In turn, PLC signaling downstream of GPCRs activates transduction channels that depolarize membrane potential (*10, 12, 54*) to activate L-type Ca^2+^ channels and ultimately trigger VSMC contraction. Our results show that stretch-induced depolarization of VSMCs is absolutely heat dependent (**Fig. 3E)**, supporting the hypothesis that transduction channels are the primary locus of thermosensitivity.

Using a combination of pharmacologic and genetic approaches, we show that TRPV1 mediates a major portion of thermo-tone in skeletal muscle both *in vitro* and *in vivo* (**Figs. 3E-K and Fig. 4)**. TRPV1 makes a greater relative contribution to tone in the 30-34°C temperature range and blockade of TRPV1 increases the T_1/2_ from ∼33°C to 36°C. Notably, stretch-induced PLC signaling lowers the temperature threshold for tone from ∼38°C to <30°C with a major contribution of TRPV1 (**Figs. 3J&K)**. Additionally, we found that TRPM4 contributes to thermo-tone in skeletal muscle arteries, particularly at temperatures greater than 34°C (**Figs. 3F,G&K**). Thus, the integration of pressure (via PLC) and temperature occurs at the transduction channels (**see Fig. 6A**). The molecular basis for thermo-tone in other tissues remains to be determined, but like skeletal muscle, it is likely that a combination of thermosensitive channels is responsible in each tissue. Indeed, previous studies have implicated TRPM4 as well as the heat-gated channels TRPV2 (*57, 58*) and ANO1 (*59*) in the myogenic tone of retinal, mesenteric, and cerebral arteries—all of these channels could confer thermo-sensitivity. On the other hand, TRP channels with limited thermo-sensitivity, such as TRPC6 (*44*) are also implicated in myogenic tone. It is possible that heat-gated channels provide the initial signaling event, which in turn, recruits thermo-insensitive channels to amplify the membrane depolarization.

**Figure 6.**
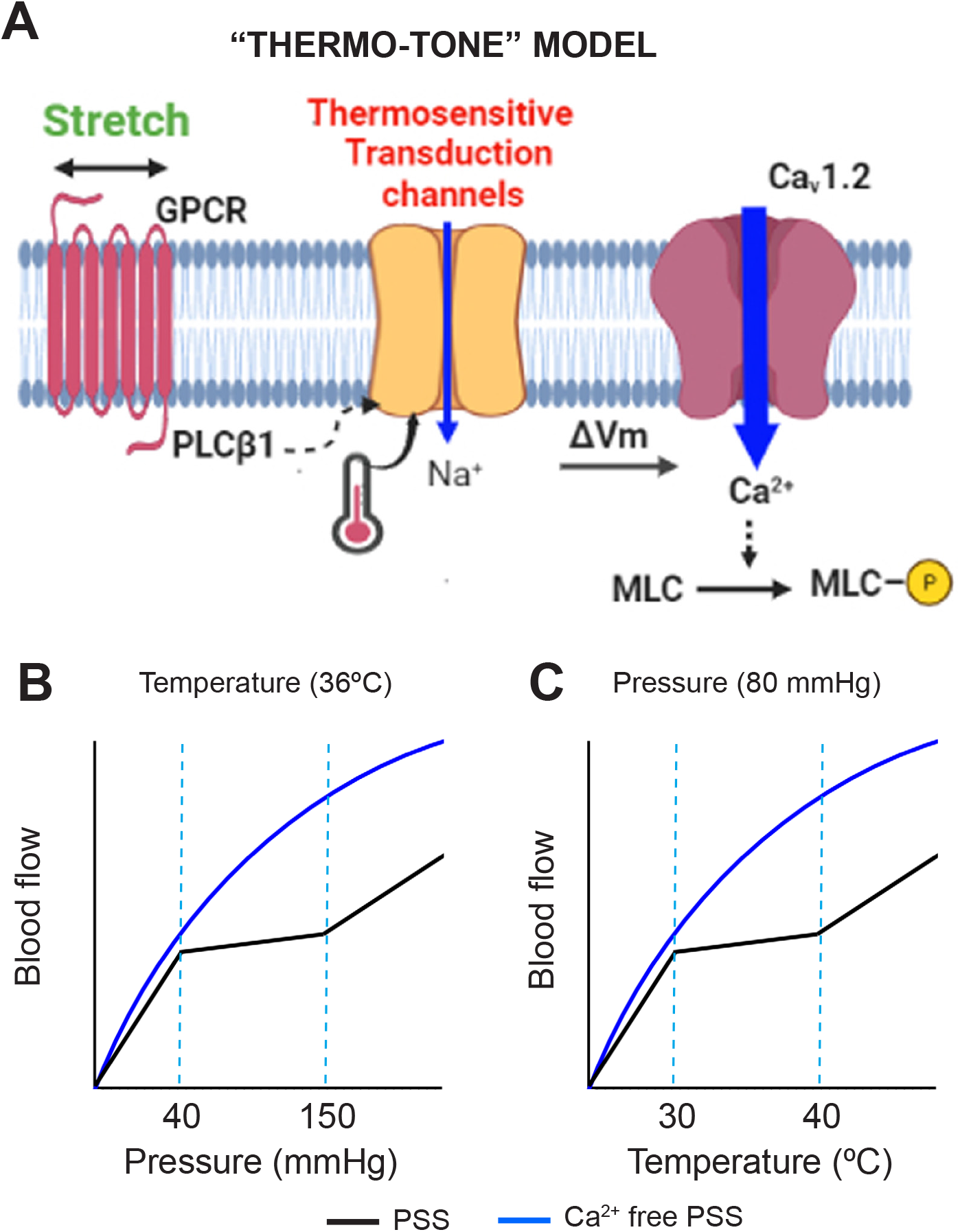
A model for temperature-dependent myogenic tone. **(A)** A proposed signaling pathway for thermo-tone. Stretch-induced PLC signaling and temperature converge to activate thermo-sensitive ion channels in VSMCs. In turn, membrane depolarization activates voltage-gated Ca^2+^ channels (Ca_V_1.2) resulting in raised myoplasmic Ca^2+^ and ultimately leading to the phosphorylation of myosin light chain necessary for contraction. **(B)** A conventional blood flow versus pressure relationship in a skeletal muscle artery (with a fixed temperature of 36°C) showing myogenic tone (black line) operating between 40 and 150 mmHg. The blue line indicates zero myogenic tone **(C)** Blood flow versus temperature in a skeletal muscle artery (with a fixed pressure of 80 mmHg) showing myogenic tone acting between 30-40°C.

We propose that thermo-tone compensates for temperature-dependent changes in the vascular conductance and local blood pressure. First, heating alters the rheological properties of blood: fluidity, RBC deformability and dispersion (*22, 23*). These changes impact flow in arterioles, capillaries, and venules where frictional forces of blood are most apparent (low Reynold’s number). In addition to rheological changes, heating also elicits vasodilation of veins, further increasing blood flow (*60*). If unchecked the effects of local heating on vascular conductance would lead to a rise in capillary pressure, resulting in tissue fluid accumulation (edema) and potential capillary damage. Indeed, we found that inhibiting myogenic tone resulted in very large increases in heat-stimulated blood flow (**Fig. 5**). Second, as blood fluidity changes with cooling or heating, the blood respectively exerts a greater or a lesser pressure on the arteriole walls. That is, the local pressure per unit flow would be either increased or reduced. A mechanism for myogenic tone solely based on pressure sensing would fail to correct for these effects to stabilize flow. These observations lead to a general principle regarding blood flow regulation: small deviations (positive or negative) from the resting tissue temperature that alter vascular conductance and blood pressure are countered by changes in thermo-tone. In our model (**Fig. 6**), myogenic tone can be represented by two independent relationships: 1) a traditional pressure-flow relationship (**Fig. 6B**), in which temperature is constant (36°C) and blood flow is stabilized over a discrete pressure range (40 to 150 mmHg), and 2) a temperature-flow relationship (**Fig. 6C**), in which pressure is fixed (80 mmHg) and blood flow is stabilized over a discrete temperature range (30-40°C). While this temperature range is specific to skeletal muscle, it can be readily adjusted to describe the thermosensitivity of other tissues. Our data therefore suggest a fundamental role for thermo-tone in the homeostatic regulation of blood flow. These findings are also relevant to the regulation of tissue perfusion during states of hypothermia or hyperthermia.

## Materials and Methods

### Ethical approval

All procedures were approved by Georgetown University, IACUC Protocol Number: 2018-0033.

### Animals

Male C57Bl6 wild-type and TRPV1-KO mice (25-30 g) were housed at 21-22°C and had *ad libitum* access to a standard laboratory chow and water.

### Ex vivo artery physiology

Mice were euthanized (CO_2_/decapitation) and the following arteries were isolated: skeletal muscle (radial artery branch, ∼60μm, or subscapular muscle branch, ∼120μm), mesenteric (3^rd^ order, ∼120μm), pial (∼80-120μm) and skin (hairy skin, and glabrous skin ∼100-150μm, see **Fig. S1**). Arteries were cannulated with glass micropipettes and secured with monofilament threads (*55*). The pipette and bathing physiological salt solution (PSS) containing (in mM): 125 NaCl, 3 KCl, 26 NaHCO_3_, 1.25 NaH_2_PO_4_, 1 MgCl_2_, 4 D-glucose, and 2 CaCl_2,_) was aerated with a gas mixture consisting of 21% O_2_, 5% CO_2_ to maintain pH (pH 7.4). To induce maximal dilation, arteries were perfused with a PSS solution containing 0 CaCl_2,_ 0.4 mM EGTA and 100 μM sodium nitroprusside (SNP). Arteries were mounted in a single vessel chamber (Living Systems Instrumentation) and placed on a heated imaging stage (Tokai Hit) to maintain bath temperature between 15-42°C, while intraluminal pressure was maintained by a Pressure Control Station (Stratagene). Arteries were viewed with a 10X objective using a Nikon TE2000 microscope and recorded by a digital camera (Retiga 3000, QImaging). Increases in intraluminal pressure or temperature were maintained for a 3-5 min duration to enable a stable vessel response. The inner arteriole diameter was measured at several locations along each arteriole using the NIH-ImageJ software’s edge-detection plug-in (Diameter) (*56*). The software automatically detects the distance between edges (by resampling a total of five times from adjacent pixels) yielding a continuous read-out ± SD of a vessel’s diameter. Myogenic tone (%) was calculated by: 100x(Diameter_ZeroCa2+_-Diameter_PSS_)/ Diameter_ZeroCa2+_. The temperature for half-maximal tone was obtained by fitting a Hill function to data from individual arteries.

### Membrane potential measurements

Sharp glass microelectrodes (50 MΩ) filled with 3 M KCl were inserted through the adventitial layer of pressurized (20 or 100 mmHg) arteries. The membrane potential was recorded using a Warner OC-725C amplifier. We restricted data analysis to impalements that produced a sharp voltage deflection and a subsequent stable potential.

### Intravital imaging

Intravital imaging was performed in anesthetized mice (urethane 1.2-1.5 g/kg/IP) as previously described (*41, 55*). Briefly, the brachial-radial artery junction was surgically exposed and visualized with a Zeiss stereomicroscope. The exposed arteries were locally perfused (using a 250 μm cannula connected to a valve-controlled gravity-fed perfusion system) with preheated PSS buffer. The surface tissue temperature was measured via a thermistor (Warner Instruments) that was positioned next to the artery. Measurements were made several minutes after temperature had stabilized.

### Laser Speckle Contrast Imaging

LSCI was performed as per intravital imaging except that the forelimb was exteriorized into a PSS-filled clear heating probe (VHP3, Moor Instruments; Devon, UK) with a temperature controller (moorVMS HEAT and LDF2). The exposed tissue was imaged with the RFLSI III system (RWD Life Science, Shenzhen, China). Resulting images were digitized and analyzed off-line.

### Systemic blood pressure recording

The experiments were performed in anesthetized mice (urethane 1.2-1.5 g/kg/IP) as previously described (*41, 55*). Briefly, the left carotid artery and the right jugular vein were cannulated with a Millar catheter (1F SPR-1000) and a polyethylene tubing (PE-10), respectively, for monitoring arterial blood pressure and for systemic (intravenous) infusion of drugs. Body temperature was maintained using a heating plate and core temperature (T_Core_) was monitored by a digital rectal thermometer and skeletal muscle (T_SkM_) temperature were monitored by a thermoprobe implanted in deep hindlimb muscle. Stock solutions of BCTC (30 mg/ml in DMSO) were dissolved in physiological saline containing 2-hydroxypropyl-Beta-cyclodextrin (25% w/v). Final DMSO concentration was <33%. Intravenous infusion of drugs was initiated only when a stable baseline of blood pressure and heart rate was present.

### Chemicals

4-(3-Chloro-2-pyridinyl)-*N*-[4-(1,1-dimethylethyl)phenyl]−1-piperazinecarboxamide (BCTC), U73122, 9-phenantrol and nifedipine were purchased from Tocris Bioscience. 4-Chloro-2-(1-naphthyloxyacetamido)benzoic acid ammonium salt (NBA) was purchased from Aobious. Stock solutions were prepared in EtOH or DMSO. Unless otherwise indicated, all other chemicals were obtained from Sigma–Aldrich.

### Statistical analysis

Data were analyzed using Prism (GraphPad Software, La Jolla, CA) and are expressed as means ± SD. Unless otherwise stated, statistical significance was evaluated using *t-* test, one-way ANOVA or one-way ANOVA with repeated measures (paired data), with treatment interactions assessed by Tukey’s *post hoc* multiple comparisons test. A *P* value of < 0.05 was considered statistically significant.

## Acknowledgments

We thank Attila Toth and Rosa Miyares for helpful suggestions and comments on the manuscript.

## Author contributions

Conceptualization: GA, TP

Methodology: TP, NS

Investigation: TP, NS, GA

Visualization: TP

Funding acquisition: GA

Project administration: GA

Supervision: GA

Writing – original draft: GA, TP

Writing – review & editing: GA, NS, TP

## Data and materials availability

All primary data are available upon request.

