## Supplementary material for "Arteries are finely tuned thermosensors regulating myogenic tone and blood flow": Fig. S1, Fig S2, Fig. S3

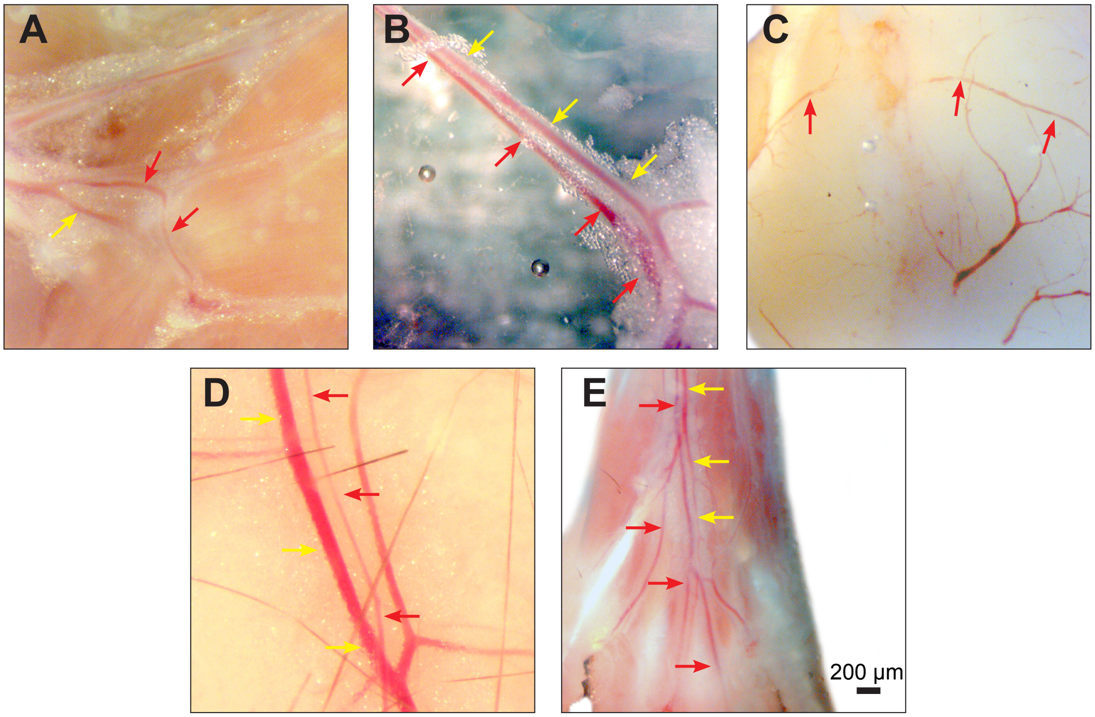


**Fig. S1. Representative images of the arteriole preparations.** A. radial muscle branch, B, mesenteric (3$^{rd}$ order), C, pial, D, hairy skin (the mid-back region) and E, digital (ventral footpad). Arteries are labeled (red arrowheads) and veins (yellow arrowheads).

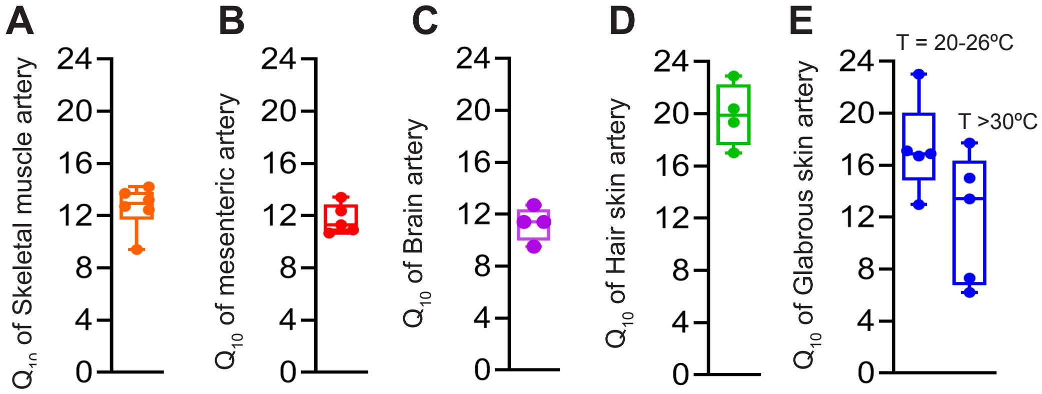


**Fig. S2. Mean $Q_{10}$ values for myogenic tone.** Data were obtained from Arrhenius plots of myogenic tone versus temperature (1000/T, $K^{-1}$) for skeletal muscle (n=6) mesenteric (n=5), pial (n=4), hairy skin (n=4) and glabrous skin (n=5).

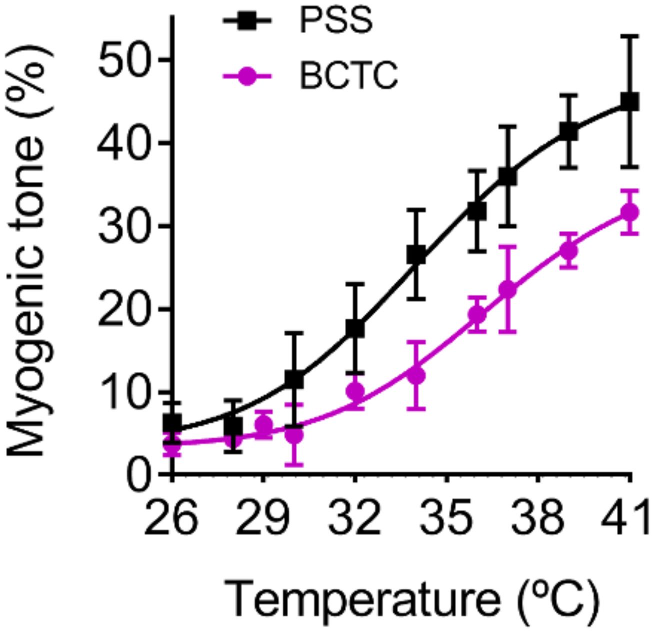


**Fig. S3. The TRPV1 inhibitor BCTC reduces the thermosensitivity of myogenic tone.** Myogenic tone versus temperature for control skeletal muscle arteries (data re-plotted from Figure 1G) and following treatment with BCTC, 3μM). BCTC increased the $T_{1/2}$ from 33.4 ± 1.8 °C (n=11) to 35.9 ± 0.6 (n=5, P=0.01).
